# Local production of reactive oxygen species drives vincristine-induced axon degeneration

**DOI:** 10.1101/2022.07.30.501173

**Authors:** Jorge Gómez-Deza, Anastasia L. Slavutsky, Matthew Nebiyou, Claire E. Le Pichon

**Author notes:** co-first authors (Jorge Gómez-Deza, Anastasia L. Slavutsky).

## Abstract

Neurological side effects arising from chemotherapy, such as severe pain and cognitive impairment, are a major concern for cancer patients. These major side effects can lead to reduction or termination of chemotherapy medication in patients, negatively impacting their prognoses. With cancer survival rates improving dramatically, addressing side effects of cancer treatment has become pressing. Here, we use iPSC-derived human neurons to investigate the molecular mechanisms that lead to neurotoxicity induced by vincristine, a common chemotherapeutic used to treat solid tumors. Our results uncover a novel mechanism by which vincristine causes a local increase in mitochondrial proteins that produce reactive oxygen species (ROS) in the axon. Vincristine triggers a cascade of axon pathology, causing mitochondrial dysfunction that leads to elevated axonal ROS levels and SARM1-dependent axon degeneration. Importantly, we show that the neurotoxic effect of increased axonal ROS can be mitigated by the small molecule mdivi-1 and antioxidants glutathione and mitoquinone, identifying a novel therapeutic avenue to treat the neurological effects of chemotherapy.

## Introduction

Vincristine is a widely used chemotherapeutic agent for treating solid tumors in children and adults [1].However, its use at high doses in patients is restricted by its potential to cause chemotherapy-induced peripheral neuropathy (CIPN) [2]. CIPN is a debilitating and sometimes irreversible condition with symptoms including chronic neuropathic pain, muscle weakness, and sensory loss [1].The emergence of CIPN symptoms often leads to a reduction or premature discontinuation of cancer treatment. To date, there is no reliable way to predict which patients may develop CIPN. Often, CIPN symptoms worsen once treatment is completed, leaving little opportunity to adjust chemotherapy frequency or dose level [3].Currently, there are limited palliative treatment options and no cure for CIPN.

Another type of neurological side effect of chemotherapeutics such as vincristine involves cognitive and memory deficits commonly referred to as chemo brain [4–6]. This suggests that central as well as peripheral neurons can be negatively affected by chemotherapy. With cancer survival rates improving significantly over the last decades, an unintended consequence is more people living with chronic pain. Thus, there is a pressing need to find effective interventions to manage vincristine-induced neurological symptoms in the growing number of affected patients [7].

Vincristine is an anti-mitotic agent that binds with high affinity to the tubulin in microtubules of the mitotic spindle and impedes cancer cell division [8]. However, vincristine also binds to microtubules of the axonal cytoskeleton, disrupting axonal transport and inducing axon degeneration by blocking microtubule polymerization [9]. In mice, vincristine treatment has been shown to cause mitochondrial transport dysfunction [10] and swelling [11], leading to increased levels of reactive oxygen species (ROS) in the soma [12]. These downstream consequences of axonal damage are potential contributors to the high incidences of neurological side effects in patients who receive treatment with vincristine.

Numerous studies have modeled vincristine-induced axon degeneration with *in vitro* assays in mouse dorsal root ganglia cultures [1, 10, 13, 14]; however, these are limited by scalability and possibly also by translational relevance to human patients. Using human neurons to find potential therapeutic interventions for neurological side effects of chemotherapy is therefore highly desirable. Inducible pluripotent stem cell-derived neuron (iPSN) technology, in addition to being human in origin, affords the production of sufficient quantities of neurons to perform large-scale analyses. Importantly, iPSNs have been used to model vincristine-induced neurotoxicity and to screen for potential therapeutic candidates [15, 16], paving the way for mechanistic studies.

In this study, we investigate the effects of vincristine on axonal health using a validated transcription-factor mediated platform to generate human neurons from iPSCs (inducible, isogenic, and integrated i^3^Neurons [17]). To identify the molecular changes that occur in human axons directly after localized exposure to vincristine, we perform mass spectrometry of i^3^Neuron axons and detect an early increase in mitochondrial and electron transport chain-related proteins. We determine that even low doses of vincristine cause mitochondrial dysfunction, increases in ROS, and mild axon degeneration. This degeneration depends on SARM1, a key executioner protein of axon degeneration [18, 19] We find that the toxic effects of vincristine can be ameliorated by mdivi-1, a small molecule reported to inhibit dynamin-related protein 1 (DRP1) and potentially interfere with mitochondrial respiratory Complex I. We demonstrate that treatment of neurons with mdivi-1 or antioxidants such as glutathione and the mitochondrial-targeted ubiquinone derivative (mitoquinone or MitoQ) reduces ROS and confers neuroprotection in the axons of human iPSC-derived neurons after vincristine treatment. We unexpectedly find that mdivi-1 mitigates vincristine-induced axon degeneration independently of DRP1, implicating the reduction of reactive oxygen species as a key target for the treatment of vincristine-induced neurotoxicity.

## Results

### Low doses of vincristine induce axon degeneration in human neurons

To investigate the effects of vincristine on human neurons, we used a transcription-factor mediated platform to generate human i^3^Neurons. i^3^ iPSCs are engineered with a doxycycline-inducible promoter to direct expression of the transcription factor NGN2 (neurogenin-2) which drives their differentiation into glutamatergic cortical-like neurons. Addition of doxycycline to the cell culture media causes iPSC differentiation into human i^3^Neurons in an efficient, scalable, and reproducible manner in 10 days [17] (Fig. 1A). Vincristine is known to cause axon degeneration *in vitro* and *in vivo* in mice [20]. To measure the effect of vincristine on human axon health, iPSCs were plated in the center of a well such that axons grew outwards, enabling the formation of a distinct axon-enriched region (Fig. 1B). Neurons were grown for 10-14 days before treatment with 5 nM vincristine, a physiologically relevant dose [21]. Axonal integrity was assessed by adapting a published method to quantify axon degeneration (Sup Fig. 1) [22]. Significant axon degeneration was observed 8 and 24 hours after vincristine treatment (Fig. 1C, D), consistent with previous results obtained in primary neurons from mouse. Thus, i^3^Neurons are a robust and scalable system to investigate the toxicity of vincristine in human neurons.

**Figure 1:**
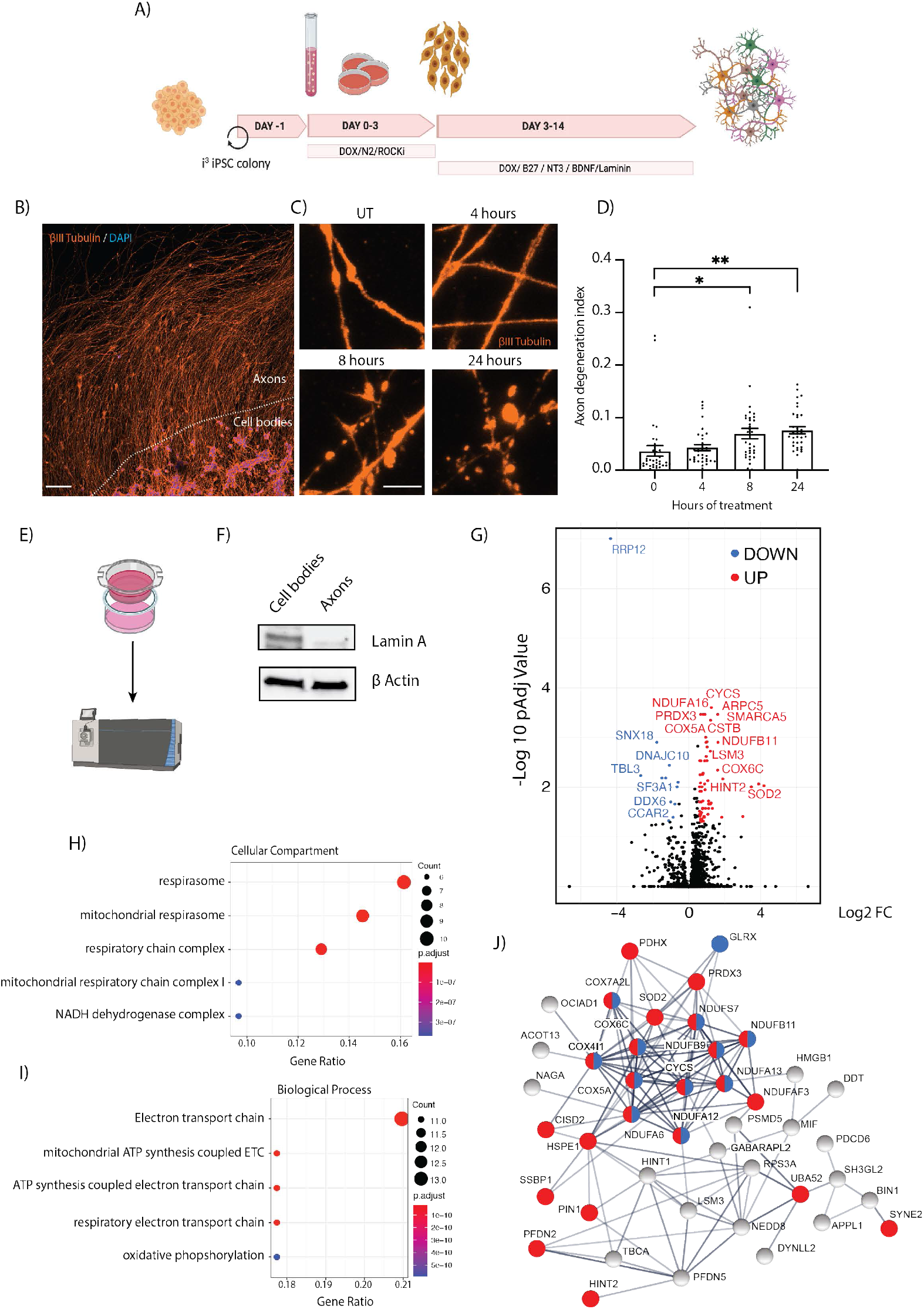
Mass spectrometry of i^3^Neuron axons reveals a vincristine-induced upregulation of mitochondrial proteins. A) Schematic representation of i^3^Neuron differentiation created using Biorender. B) Representative image of i^3^Neuron cell bodies plated in the center of an 8-well slide to allow axons to grow outwards. Immunostaining for βIII tubulin (orange), DAPI (blue). 20X magnification. A = Axon area (Scale bar =100 μm). C) Representative images of axon degeneration upon addition of 5 nM vincristine for 0, 4, 8, or 24 hours. Immunostaining for βIII tubulin (orange), DAPI (blue) (Scale bar =5 μm). D) Axon degeneration index (ADI) quantification for i^3^Neuron-treated axons with 5 nM vincristine. N=5 independent differentiations. One-way ANOVA, Bonferroni correction (p<0.05 *, p<0.01 **). E) Schematic representation of Boyden chambers for axonal separation and experimental procedure (created using Biorender). F) Western blotting showing efficient isolation of axons from cell bodies. The nuclear envelope protein Lamin A is absent from the axon fraction. Uncropped western blots presented in Sup Fig 5. G) Volcano plot showing the significant proteins up (red) and down (blue) regulated in i^3^Neuron axons after 5 nM vincristine treatment for 4 hours (>2 unique peptides detected, fold change (FC) >1.5, adjusted P-value <0.05.) H) Dot plot showing top 5 enriched cellular compartment gene ontology analysis categories for axon-enriched proteins after 5 nM vincristine treatment ranked by gene ratio. I) Dot plot showing top 5 enriched biological process gene ontology analysis categories for axon-enriched proteins after 5 nM vincristine treatment ranked by gene ratio. J) Protein-protein interaction network for proteins upregulated in the axon after vincristine treatment. Mitochondrial proteins are highlighted in red and electron transport chain proteins in blue. Proteins involved in the catalysis of ROS were also elevated (Superoxide dismutase 2 (SOD2), Thioredoxin-dependent peroxide reductase (PRDX3), Glutaredoxin-1 (GLRX) and Thioredoxin Domain Containing 17 (TXNDC17).

### Mass spectrometry of i^3^Neuron axons reveals vincristine-induced mitochondrial deficits

We predicted that vincristine application would cause acute protein changes in axons. Therefore, we locally applied vincristine just to the axons and selectively purified this cellular compartment for unbiased mass spectrometry analyses. Neurons were plated in Boyden chambers (Fig. 1E) which allow axons to grow through a porous membrane and be cleanly isolated from somas, as assessed by the absence of the nuclear protein Lamin A in the axonal fraction (Fig. 1F). Using this platform, we compared control axons treated with DMSO with those selectively treated with 5 nM vincristine. After 4 hours, the axonal and cell body compartments were harvested and subjected to label-free mass spectrometry. In total, 3,441 proteins were identified, with 766 enriched in soma and 224 in axons (Sup Fig. 2A, Sup File 1). Gene ontology analysis of proteins significantly enriched in the cell body highlighted proteins involved in RNA catabolic process, translation initiation, and RNA splicing (Sup Fig. 2B). Conversely, proteins involved in axonogenesis, synapse organization, and axon guidance were enriched in the axonal fraction (Sup Fig. 2C). These results validated our approach to efficiently isolate axons from the cell bodies and provide a rich dataset to investigate protein level changes during CIPN.

When comparing vincristine to DMSO-treated axons, 63 enriched and 12 depleted proteins were identified (>2 unique peptides detected, fold change >1.5, adjusted P-value <0.05, Fig. 1G, Sup File 1). Notably, gene ontology analysis revealed that a significant fraction of the upregulated proteins was mitochondrial (23/63; 36%) (Fig. 1H) and involved in ATP synthesis or oxidative phosphorylation (Fig. 1I). String analysis was performed to create a protein-protein interaction network of vincristine-induced upregulated proteins. This analysis showed that many of the enriched mitochondrial proteins are known to interact (Fig. 1J, red). Interestingly, we observed largely connected cluster that is associated with the electron transport chain, an intensive site of ROS production [23] (Fig. 1J, blue). This led us to hypothesize that vincristine causes axon degeneration by impairing axonal mitochondrial function and inducing oxidative stress through the overproduction of reactive oxygen species [23].

### Vincristine causes mitochondrial dysfunction and increased ROS in i^3^Neuron axons

We next addressed the effect of vincristine on mitochondrial function. Mitochondrial health can be evaluated by measuring cellular oxygen consumption rates using a Seahorse metabolic flux assay. Exposing neurons for 4, 8, and 24 hours to 5 nM vincristine caused basal respiration levels to drop significantly for the next 24 hours (Fig. 2A, B). These data demonstrate that a major effect of vincristine is impairment of normal mitochondrial function in neurons.

**Figure 2:**
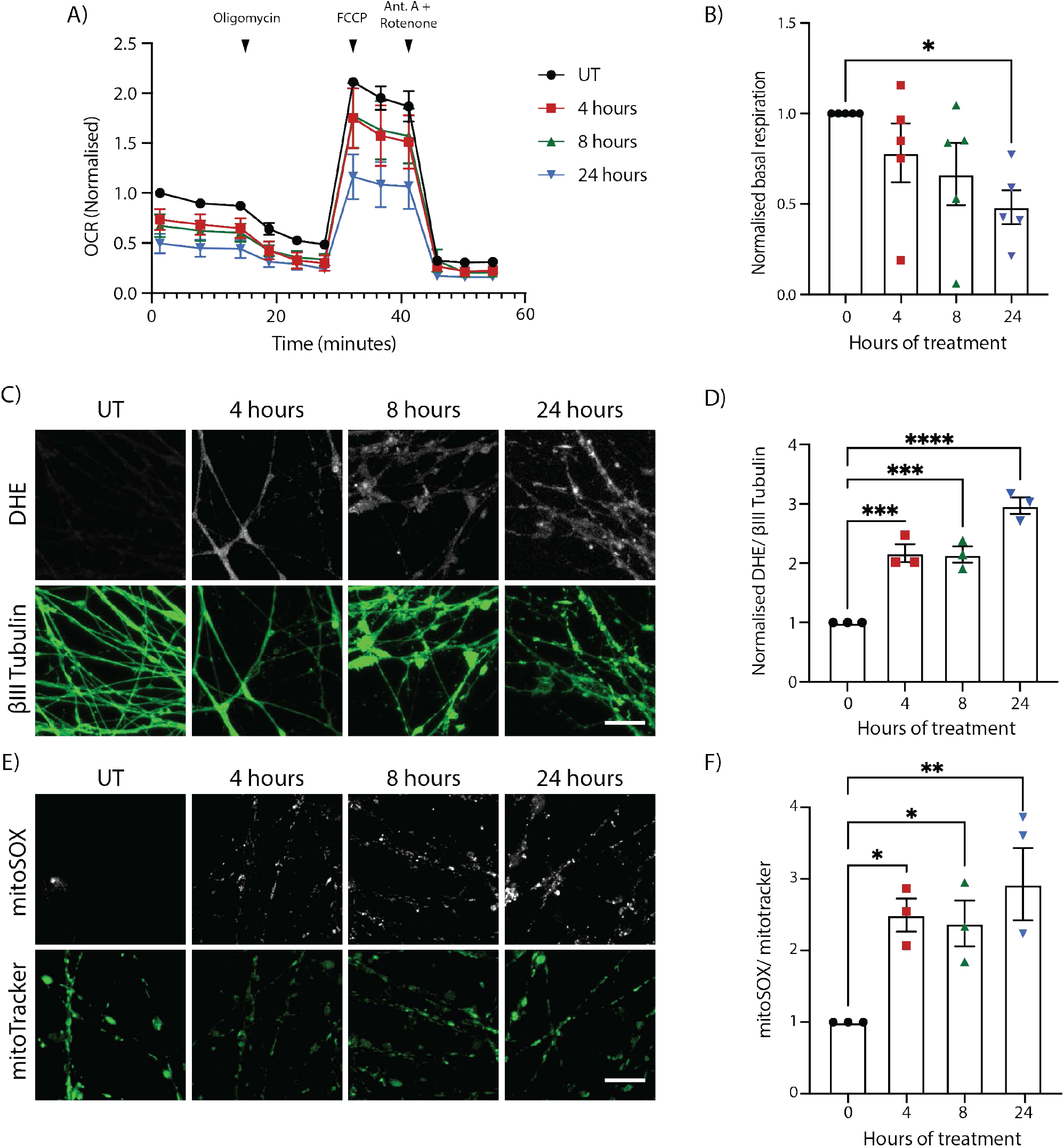
Vincristine causes mitochondrial dysfunction and increased ROS in i^3^Neurons. A) Seahorse oxygen consumption rate (OCR) analysis of i^3^Neurons untreated (UT) or treated for 4, 8, and 24 hours with 5 nM vincristine. Results normalized to UT i^3^Neurons. Results are represented as mean ± SEM. N=5 independent differentiations, 3 wells measured per condition. Oligomycin (2 μM), FCCP (1 μM) and antimycin A/rotenone (500 nM) were added as indicated. B) Basal OCR levels of i^3^Neurons UT and treated for 4, 8, and 24 hours with 5 nM vincristine. Results normalized to UT i^3^Neurons. Results are represented as mean ± SEM. N=5 independent differentiations, 3 wells measured per condition. One-way ANOVA, Bonferroni correction (p<0.05 *). C) Representative images of 5 nM vincristine-treated i^3^Neuron axons for ROS quantification. Staining with DHE (white) and βIII tubulin (green). Scale bar = 20 μm. D) Quantification of DHE fluorescence in 5nM vincristine-treated i^3^Neuron axons. Results normalized to untreated axons. Results are represented as mean ± SEM. N=3 independent differentiations, 5-8 images per differentiation. One-way ANOVA, Bonferroni correction (p<0.005 ***). E) Representative images of 5 nM vincristine-treated i^3^Neuron axons for ROS quantification. Staining with mitoSOX (white) and mitoTracker green (green). Scale bar = 20 μm. F) Quantification of mitoSOX fluorescence in 5 nM vincristine-treated i^3^Neuron axons. Results normalized to UT axons. Results are represented as mean ± SEM. N=3 independent differentiations, 5-8 images per differentiation. One-way ANOVA, Bonferroni correction (p<0.05 *, p<0.01 **).

Mitochondrial dysfunction is known to result in an increase in reactive oxygen species [23]. Additionally, our mass spectrometry results showed that proteins involved in oxidative phosphorylation as well as those involved in the catalysis of ROS (SOD2, PRDX3, GLRX and TXNDC17) were increased after axonal exposure to vincristine. We hypothesized that vincristine causes a local increase in the levels of ROS in human axons leading to axon degeneration. To measure the levels of ROS in axons, neurons were incubated with dihydroethidium (DHE), a cell-penetrable fluorescent indicator of reactive oxygen species in the cytoplasm [24]. Treatment of i^3^Neurons with vincristine resulted in a significant increase in axonal ROS levels at 4, 8, and 24 hours after treatment (Fig. 2C, D).

Mitochondrial dysfunction and cellular stress can also lead to an increase in mitochondrial superoxide species. We hypothesized that a significant proportion of the observed increase in total axonal ROS levels would be accounted for by mitochondrial ROS. Mitochondrial ROS levels in the axon were indeed increased 4, 8, and 24 hours after exposure to vincristine as measured by mitoSOX (Fig. 2E, F). Taken together, these results demonstrate that vincristine induces mitochondrial dysfunction and leads to an increase in total and mitochondrial ROS levels in axons.

### Mdivi-1 inhibits vincristine-induced axon degeneration

We hypothesized that inhibiting ROS generation may delay axon degeneration. To test this, vincristine-treated neurons were exposed to the small molecule mdivi-1. Mdivi-1 is often described as a DRP1 inhibitor, and has been utilized in the context of neuronal injury to attenuate traumatic brain injury induced-cell death [25], protect against glutamate excitotoxicity and oxygen-glucose deprivation [26], and reduce cerebral damage caused by ischemia/reperfusion injury [27]. However, it has also been shown to reduce ROS production in neurons [28]. Through this mechanism, mdivi-1 has conferred neuroprotection by suppressing the mitochondrial apoptosis pathway and decreasing ER stress and ROS-mediated oxidative stress [29].

When we treated neurons with 50 μM mdivi-1 in addition to 5 nM vincristine, axon degeneration was significantly reduced by two-fold after 24 hours compared to neurons treated only with vincristine (Fig. 3A, B). We next asked whether mdivi-1 acts locally on the axon, or whether application to the whole cell is required for its protective effect. Neurons were grown in microfluidic chambers to physically isolate axons from cell bodies (Fig. 3C). Vincristine, mdivi-1, or their combination was added to the axonal compartment for 24 hours. We observed no degeneration in axons treated with mdivi-1 alone (Fig. 3D, E). However, local treatment of axons with mdivi-1 significantly reduced vincristine-induced axonal degeneration (Fig. 3D, E). Together, these data reveal that vincristine acts locally to induce axonal degeneration and that axonal application of mdivi-1 can delay this degeneration.

**Figure 3:**
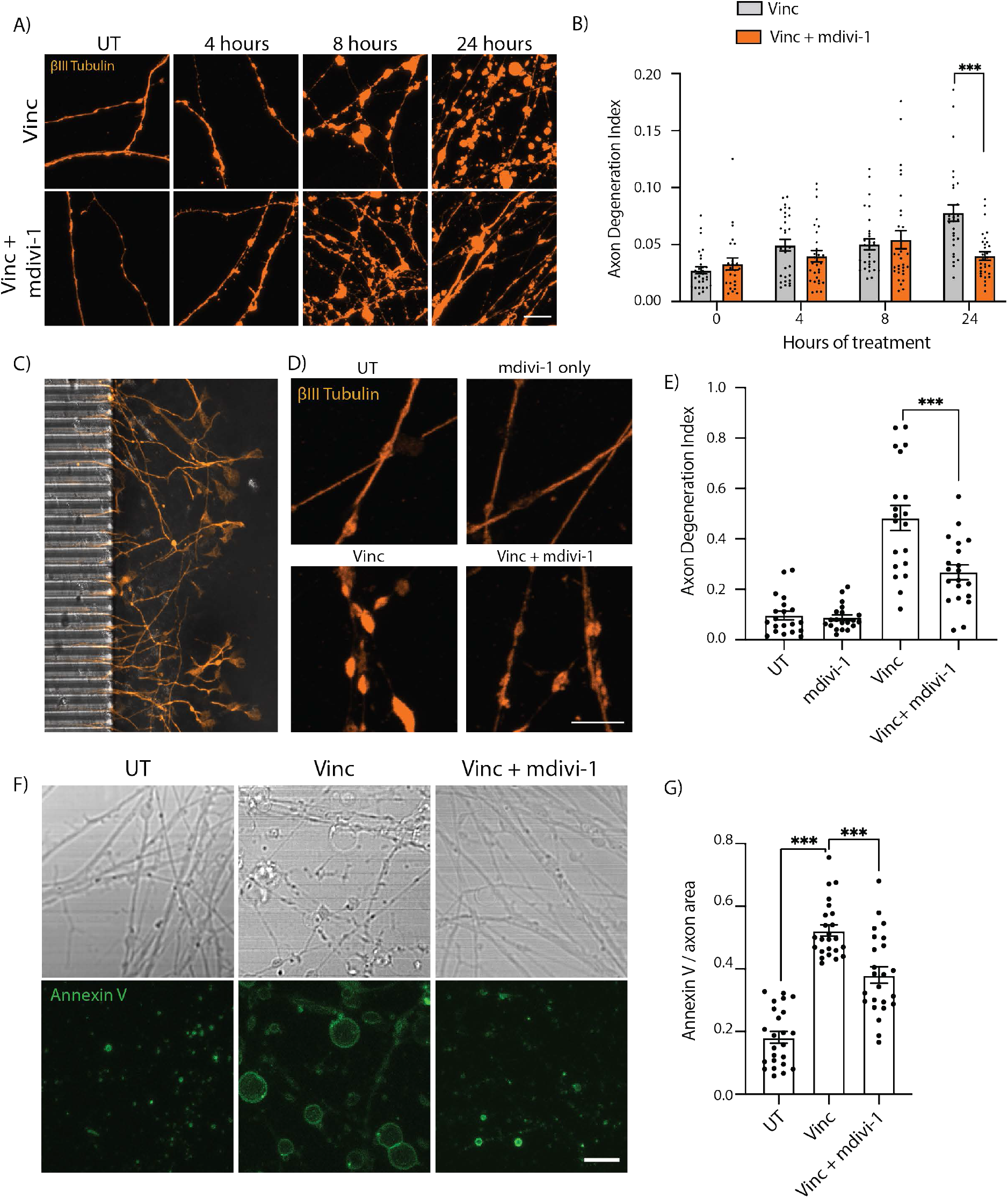
Mdivi-1 delays vincristine-induced axonal degeneration. A) Representative images of i^3^Neuron axons treated with 5 nM vincristine or 5 nM vincristine + 50 μM mdivi-1. Immunostaining for βIII tubulin (orange). Scale bar =10 μm. B) Axon degeneration index for the experiments shown in A. Results are represented as mean ± SEM. N=3 independent differentiations, 10 images per differentiation. Twoway ANOVA, Bonferroni correction (p<0.05 *). C) Image of the axon compartment of a microfluidic chamber used for isolation of axons and cell bodies, with the microgrooves for axons visible on the left. D) Representative images of UT, 50 μM mdivi-1, 5 nM vincristine, and 5nM vincristine + 50 μM mdivi-1 treated i^3^Neuron axons for 24 hours in the microfluidic axonal compartment. Immunostaining for βIII tubulin (orange). Scale bar =10 μm. E) Axon degeneration index of the experiments shown in D. Results are represented as mean ± SEM. N=3 independent differentiations, 5-8 images per differentiation. Oneway ANOVA, Bonferroni correction (p<0.005 ***). F) Representative brightfield and Annexin V stain images of UT, 5 nM vincristine and 5 nM vincristine + 50 μM mdivi-1 treated i^3^Neuron axons for 24 hours. Scale bar =10 μm. G) Quantification of Annexin V staining of UT, 5 nM vincristine and 5nM vincristine + 50 μM mdivi-1 treated i^3^Neuron axons for 24 hours. N=3 individual differentiations. N=3 independent differentiations, 5-8 images per differentiation. One-way ANOVA, Bonferroni correction (p<0.005 ***).

We also measured axon degeneration by another method, using a fluorescently-labeled Annexin V conjugate in live neurons. In apoptotic cells, Annexin V accumulates in the outer plasma membrane due to its high affinity for phosphatidylserine, which undergoes externalization during apoptosis [30, 31]. An increase in the levels of Annexin V fluorescence correlates with apoptotic cell death, or in this case, externalization of axonal membrane and axon degeneration. Center-plated neurons were treated with vincristine and mdivi-1 for 24 hours and then stained with Annexin V. Neurons treated with vincristine had significantly greater levels of Annexin V staining than untreated neurons (Fig. 3F and G). Mdivi-1 significantly reduced Annexin V staining (Fig. 3F, G). These data demonstrate that axonal exposure to vincristine is sufficient to cause axon degeneration that can be mitigated by mdivi-1.

### Preventing mitochondrial fission does not delay vincristine-induced axon degeneration

Because mdivi-1 has been used as an inhibitor of DRP1 and thus of mitochondrial fission [32], we asked whether inhibiting mitochondrial fission would reduce vincristine-induced axon degeneration. To knock down DRP1, I^3^ iPSCs containing dCas9 gene silencing machinery were transduced with gRNAs to DNM1L (which encodes DRP1) or controls [33]. Using this approach, an 80 percent reduction in the DRP1 transcript levels was measured by qPCR (Fig. 4A). DRP1 knockdown was functionally confirmed in mature neurons by measuring the mitochondrial size in control and DRP1 knockdown (KD) neurons (Fig. 4B, C). As expected, DRP1 KD neurons had significantly larger mitochondria (~60% increase) (Fig. 4B, C), validating that the knockdown of DRP1 in i^3^Neurons impairs mitochondrial fission.

**Figure 4:**
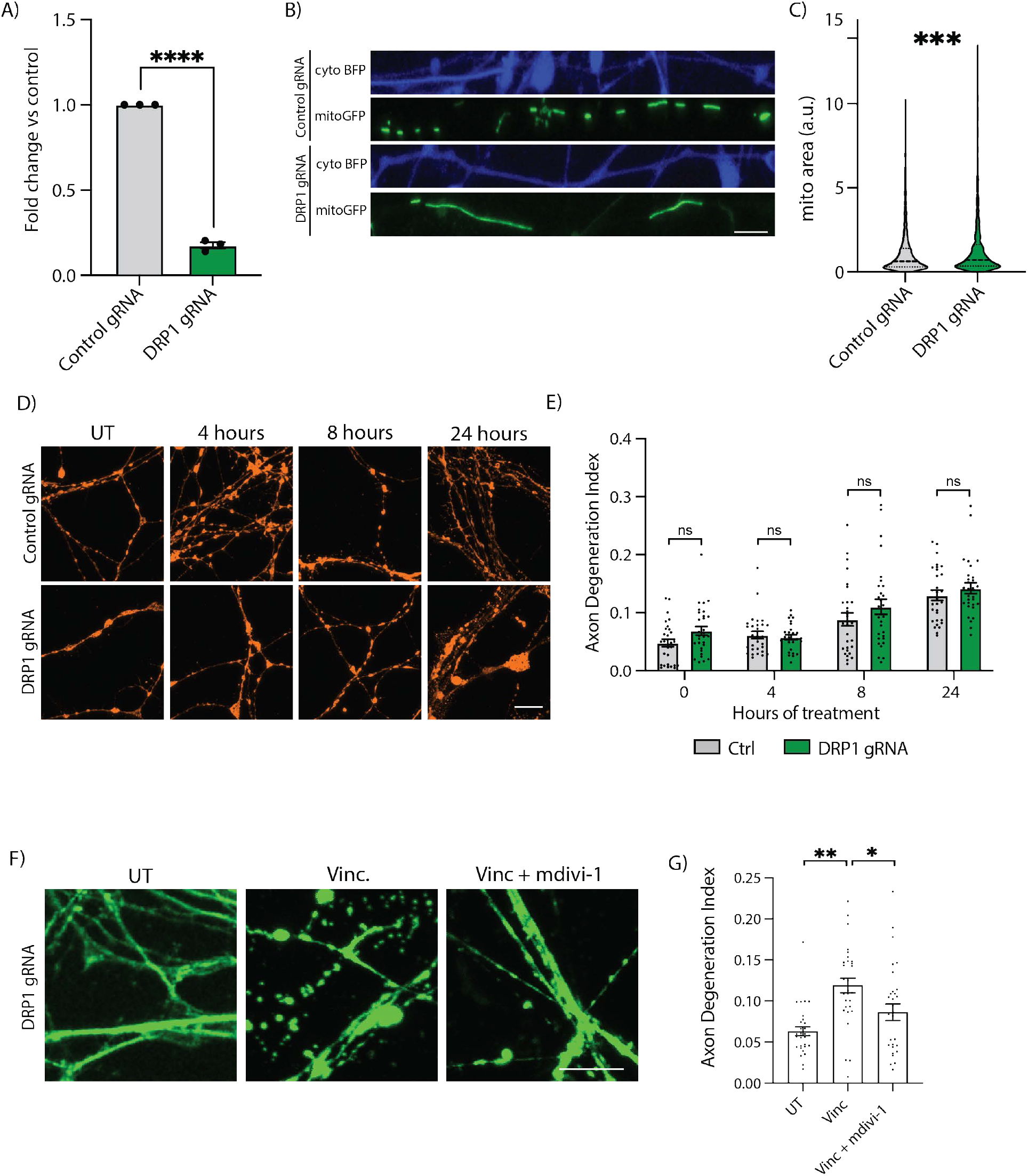
Mdivi-1 delays axon degeneration via a DRP1-independent mechanism. A) Validation of DRP1 gene knockdown (KD) using qPCR in CRISPRi-i^3^ iPSCs. Results are represented as mean ± SEM. N=3. Unpaired t-test. (p<0.0001 ****). B) Representative images of axonal mitochondria in control and DRP1 KD i^3^Neurons. Axon (cytoplasmic blue fluorescent protein (BFP)) and mitochondria (mitoGFP). Scale bar = 10 μm. C) Quantification of mitochondrial area in control and DRP1 KD i^3^Neurons. Results are represented as mean ± SEM. Over 2,000 mitochondrial particles quantified from 3 independent differentiations. Unpaired T-test (p<0.005 ***). D) Representative images of control and DRP1 KD i^3^Neuron axons for 4, 8, and 24 hours with 5 nM vincristine for axon degeneration quantification. Immunostaining for βIII tubulin (orange). Scale bar = 10 μm. E) Axon degeneration index of the experiments shown in D. Results are represented as mean ± SEM. N=3 independent differentiations, 5-8 images per differentiation. Twoway ANOVA, Bonferroni correction. No significant differences. F) Representative images of control and DRP1 KD i^3^Neuron axons UT or treated for 24 hours with 5 nM vincristine and 5 nM vincristine + 50 μM mdivi-1. Scale bar = 10 μm. G) Axon degeneration index of the experiments shown in G. Results are represented as mean ± SEM. N=3 independent differentiations, 8-10 images per differentiation. Twoway ANOVA, Bonferroni correction. (p<0.05 *, p<0.001 **).

We next treated control and DRP1 KD neurons with vincristine. Inhibiting mitochondrial fission via DRP1 knockdown failed to delay or reduce vincristine-induced axon degeneration (Fig. 4D, E). Moreover, DRP1 phosphorylation (p-S616), a marker of increased mitochondrial fission [34], was not increased after vincristine treatment (Sup. Fig. 3A, B and Sup. Fig. 5). Vincristine treatment did not increase the number of axonal mitochondrial particles (Sup. Fig. 3C, D) as would be expected if it induced mitochondrial fission. Finally, treatment of DRP1 KD neurons with mdivi-1 resulted in a significant reduction in vincristine-induced axon degeneration after 24 hours (Fig 4F, G). Together, our results demonstrate that vincristine does not induce mitochondrial fission in i^3^Neurons and inhibiting mitochondrial fission does not delay vincristine-induced axonal degeneration. We therefore conclude that mdivi-1 delays axonal degeneration via a DRP1-independent mechanism.

### Mdivi-1 reduces vincristine-induced axonal ROS levels

Mdivi-1 has recently been shown to act not by inhibiting DRP1, but instead by inhibiting ROS production in mitochondria [28, 35–37]. We therefore evaluated whether mdivi-1 treatment resulted in a decrease in axonal ROS levels. I^3^Neurons were treated with vincristine alone or concurrently with mdivi-1 for 4, 8, and 24 hours, and at each timepoint, concurrent treatment with vincristine and mdivi-1 resulted in a significant decrease in axonal ROS levels measured by dihydroethidium (DHE) compared to treatment with vincristine alone (Fig. 5A, B). Additionally, mdivi-1 significantly reduced the levels of mitochondrial ROS 8 and 24 hours after vincristine treatment (Fig. 5C, D). Here we demonstrate that mdivi-1 delays vincristine-induced axonal degeneration by reducing the levels of ROS and not by inhibiting mitochondrial fission.

**Figure 5:**
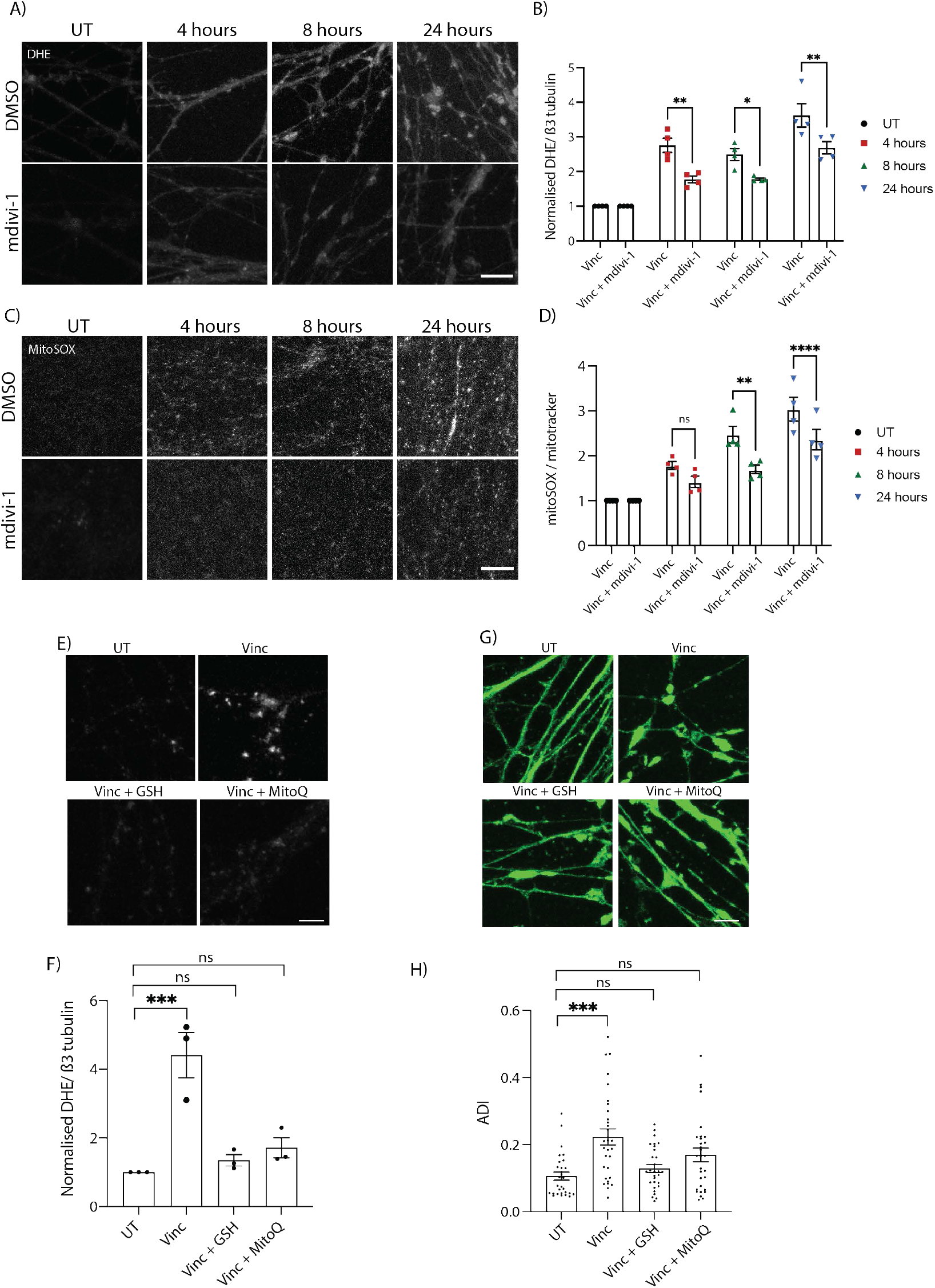
Local reduction of ROS in the axon delays vincristine-induced axon degeneration. A) Representative images of 5 nM vincristine-treated and 5 nM vincristine + 50 μM mdivi-1 treated i^3^Neuron axons. Staining with DHE (white). Scale bar = 20 μm. B) Quantification of DHE fluorescence in 5 nM vincristine and 5 nM vincristine + 50 μM mdivi-1 treated i^3^Neuron axons shown in A. Results normalized to untreated axons. Results are represented as mean ± SEM. N=4 independent differentiations, 5-8 images per differentiation. Two-way ANOVA, Bonferroni correction (p<0.05 *, p<0.001 **, p<0.005 ***). C) Representative images of 5 nM vincristine-treated and 5 nM vincristine + 50 μM mdivi-1-treated i^3^Neuron axons ROS quantification. Staining with mitoSOX (white). Scale bar = 20 μm. D) Quantification of mitoSOX fluorescence in 5 nM vincristine and 5 nM vincristine + 50 μM mdivi-1-treated i^3^Neuron axons shown in C. Results normalized to untreated axons. Results are represented as mean ± SEM. N=4 independent differentiations, 5-8 images per differentiation. Two-way ANOVA, Bonferroni correction (p<0.05 *, p<0.001 **). E) Representative images of UT, vincristine, vincristine + 5 mM GSH and vincristine + 5 μM MitoQ treated i^3^Neuron axons for 24 hours. Staining with DHE (white). Scale bar = 10 μm. F) Quantification of DHE fluorescence shown in G. Results normalized to untreated axons. Results are represented as mean ± SEM. N=3 independent differentiations, 810 images per differentiation. Two-way ANOVA, Bonferroni correction (p<0.05 *, p<0.001 **, p<0.005 ***). G) Representative images of UT, vincristine, vincristine + 5 mM GSH and vincristine + 5 μM MitoQ treated i^3^Neuron axons for 24 hours. Scale bar = 10 μm. H) Axon degeneration index of the experiments shown in E. Results are represented as mean ± SEM. N=3 independent differentiations, 8-10 images per differentiation. Twoway ANOVA, Bonferroni correction. (p<0.005 ***).

To further test whether reducing ROS prevented axon degeneration, neurons were treated with two potent antioxidant agents that have been shown to inhibit ROS after neuronal stress; glutathione (GSH) [38] and mitoquinone (MitoQ) [12, 39]. As expected, treatment with GSH or MitoQ almost completely abolished vincristine-induced ROS generation in the axons after 24 hours (Fig. 5E, F). Furthermore, GSH and to a lesser extent MitoQ were able to delay vincristine-induced axon degeneration (Fig. 5G, H). These results demonstrate that inhibition of ROS in axons can reduce vincristine-induced axon degeneration.

### Mitochondrial dysfunction and ROS production lies upstream of SARM1

Mitochondrial dysfunction and an increase in ROS levels has been shown to lie upstream of sterile α and TIR motif containing 1 (SARM1) to cause axon degeneration in the cell bodies of mouse neurons in vitro [18]. To confirm this in human axons in response to vincristine, we generated SARM1 KO iPSCs using CRISPR/Cas9 gene editing (Sup. Fig. 4). Wildtype and SARM1 KO neurons were treated with 5 nM vincristine for 4, 8, and 24 hours (Fig. 6A, B). Deletion of SARM1 significantly reduced axon degeneration approximately two- and four-fold 8 and 24 hours after vincristine treatment, respectively, compared to wildtype neurons. To confirm that SARM1 was acting downstream of the ROS increase caused by vincristine, we measured ROS levels in wildtype and SARM1 KO neurons. We found no significant difference in the levels of total axonal ROS (Fig. 6C, D) or mitochondrial ROS levels (Fig. 6E, F) between wildtype and SARM1 KO neurons in response to vincristine treatment. Taken together, our results show that mitochondrial dysfunction and an increase in ROS levels in the axon lie upstream of SARM1 activation and axon degeneration in human neurons.

**Figure 6:**
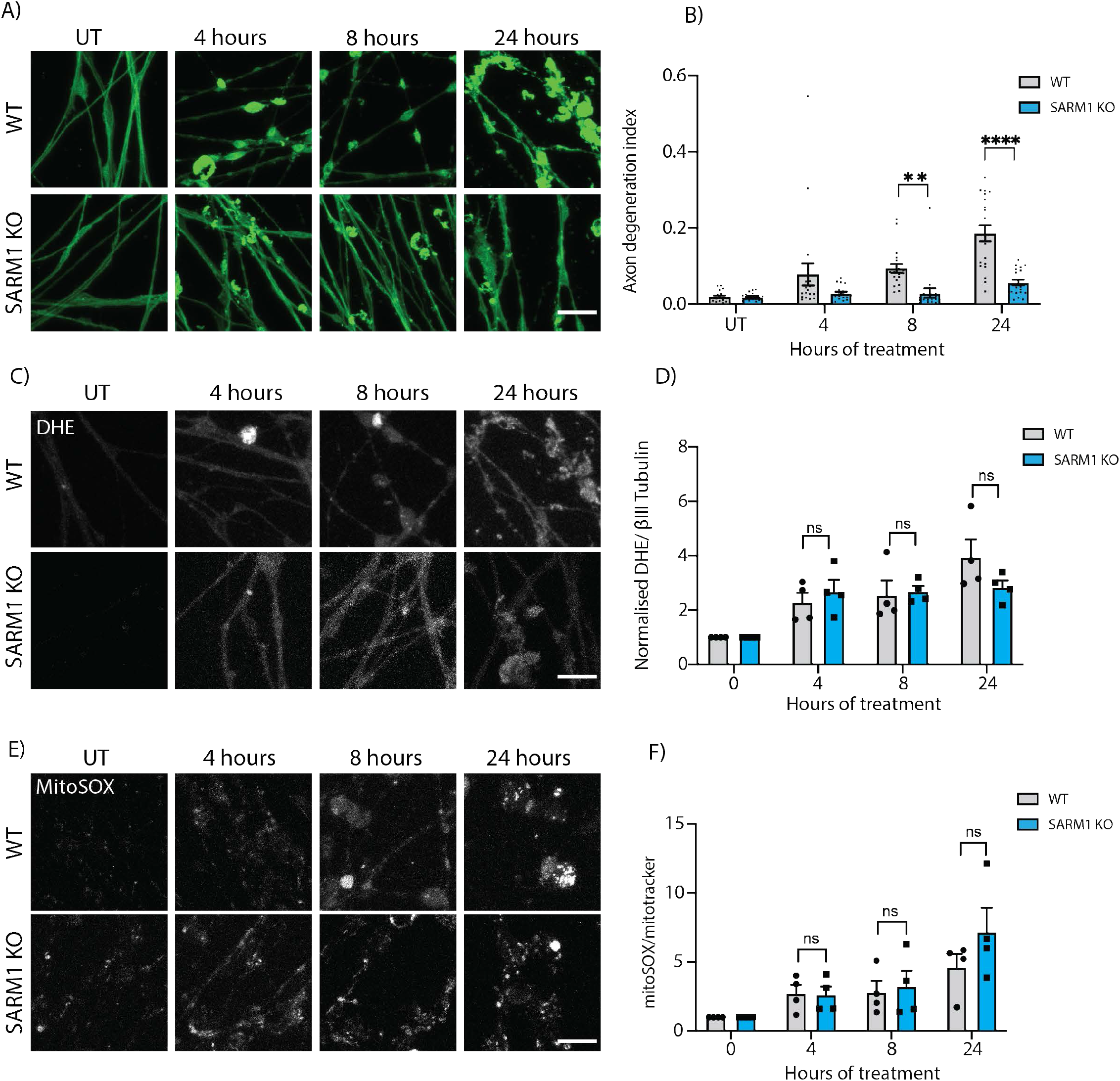
SARM1 knockout prevents axon degeneration and does not prevent axonal and mitochondrial ROS production after vincristine treatment. A) Representative images of 5 nM vincristine-treated WT and SARM1 KO i^3^Neuron axons. Immunostaining for βIII tubulin (green). Scale bar =20 μm. B) Axonal degeneration index of the experiments shown in A. Vincristine treated WT and SARM1 KO i^3^Neuron axons. Results are represented as mean ± SEM. N=4 independent differentiations, 5 images per differentiation. Two-way ANOVA, Bonferroni correction (p<0.01 **, p<0.001 ***). C) Representative images of 5 nM vincristine-treated WT and SARM1 KO neurons. Staining with DHE (white). Scale bar = 20 μm. D) Quantification of DHE fluorescence in 5 nM vincristine WT and SARM1 KO i^3^Neuron axons shown in C. Results normalized to untreated axons. Results are represented as mean ± SEM. N=4 independent differentiations, 5-10 images per differentiation. Twoway ANOVA, Bonferroni correction (ns = not significant). E) Representative images of 5 nM vincristine-treated WT and SARM1 KO i^3^Neuron axons ROS quantification. Staining with mitoSOX (white). Scale bar = 20 μm. F) Quantification of mitoSOX fluorescence in 5 nM vincristine WT and SARM1 KO i^3^Neuron axons shown in E. Results normalized to untreated axons. Results are represented as mean ± SEM. N=4 independent differentiations, 5-10 images per differentiation. Two-way ANOVA, Bonferroni correction (ns = not significant).

**Figure 7:**
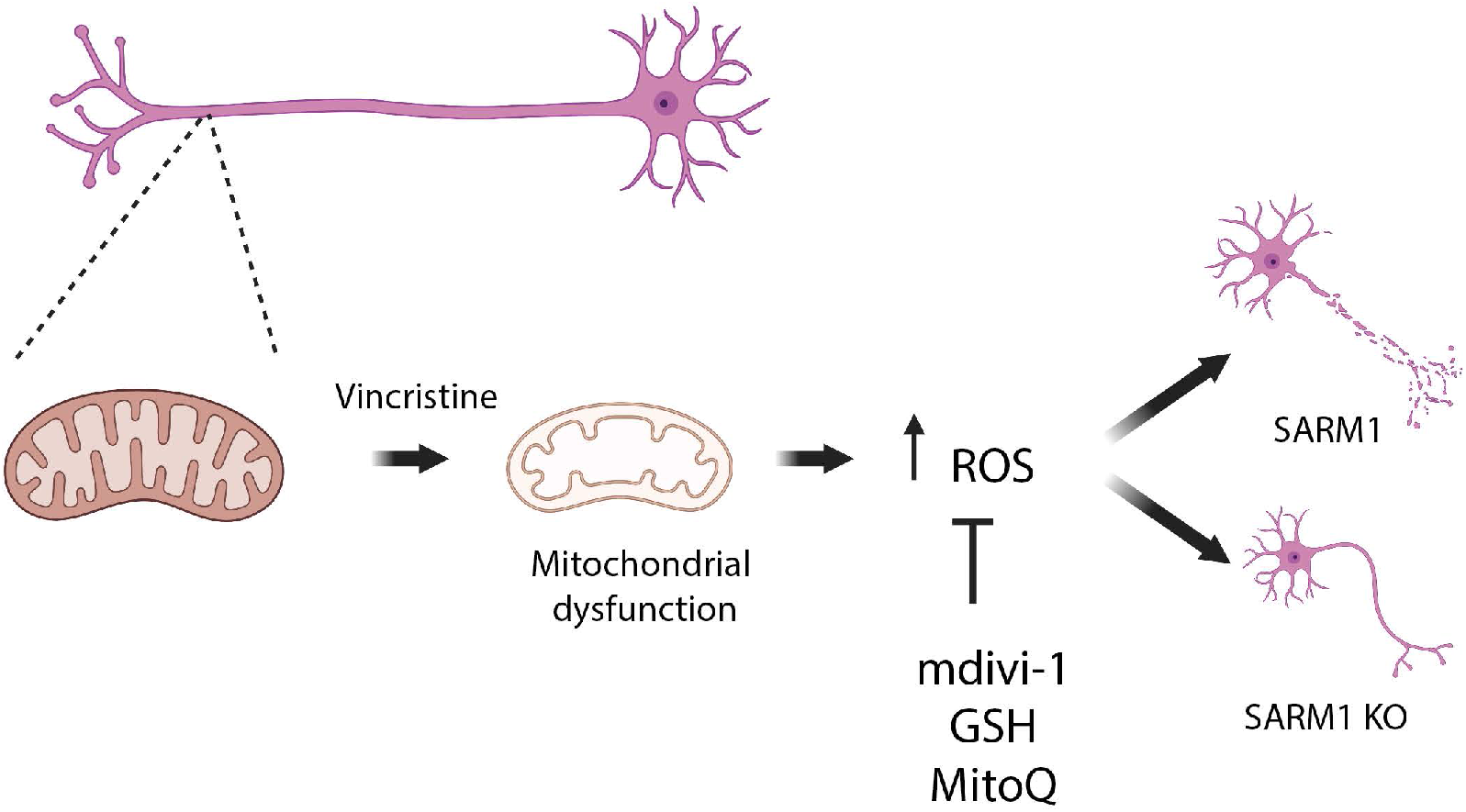
Proposed model for vincristine-induced axon degeneration in human neurons.

## Discussion

In the present study, we validate i^3^Neurons as a highly translatable model for studying vincristine-induced axon degeneration. We demonstrate that an increase in mitochondrial proteins involved in oxidative phosphorylation and increase of ROS in the axon is an early molecular event in response to vincristine. We uncover a novel mechanism by which vincristine causes mild mitochondrial dysfunction, leading to increased levels of reactive oxygen species in the axon, and to SARM1-dependent axon degeneration. Additionally, we show that mdivi-1 and other antioxidants reduce vincristine-induced axon degeneration by decreasing levels of ROS in the axon.

Previous studies have used iPSC-derived neurons to model the effect of vincristine on neuronal survival [15, 40–45]. However, these studies typically used much higher doses of vincristine (50 nM – 1 μM) than levels that have been measured in cerebral spinal fluid of human patients after vincristine treatment (~0.1 – 2 nM) [21, 46]. This discrepancy of 25-500-fold in vincristine dose level calls into question the ability of previous studies to accurately model the effects of vincristine on neuronal health. In our study, we used 5 nM vincristine which is much closer to the actual levels measured in patients undergoing chemotherapy.

Vincristine is known to cause mitochondrial depolarization [47], but the mechanism by which this happens remains poorly understood. We found that vincristine acts locally to increase the levels of axonal mitochondrial proteins that constitute Complex I of the electron transport chain involved in oxidative phosphorylation and catalysis of reactive oxygen species. This correlated well with our observed increase in the levels of axonal ROS. However, the rates of oxidative phosphorylation were unchanged 4 hours after vincristine treatment as measured by the seahorse assay. We hypothesize that the enrichment in mitochondrial proteins may be an adaptive response to mitochondrial dysfunction caused by vincristine in the axon, since increased assembly of complex I would be predicted to result in a reduction of reactive oxygen species [48]. Elevated ROS levels have been reported in the spinal cord of mice treated with vincristine [12] and in the brains of patients that undergo chemotherapy [4]. We observed increased levels of total and mitochondrial ROS in the axon as early as 4 hours after vincristine treatment, in agreement with these studies. Interestingly, since the changes in protein levels happened on the scale of just a few hours after the addition of vincristine, it is possible that they arise from local regulation in the axon (such as protein stabilization or local translation). This merits further investigation in future.

We also demonstrate that mdivi-1 reduces vincristine-induced axon degeneration by locally reducing the levels of ROS in the axon. Although it was initially discovered as an inhibitor of mitochondrial fission [32], mdivi-1 has since been shown to mitigate the production of ROS, possibly through the inhibition of Complex I of the ETC [28]. In our study, we observed no evidence of mitochondrial fission after exposure to vincristine. Furthermore, knockdown of DRP1 did not reduce vincristine-induced axon degeneration, and treatment of DRP1-deficient neurons with mdivi-1 attenuated vincristine-induced axon degeneration. We therefore concluded that mdivi-1 protects axons from vincristine-induced degeneration by reducing ROS and not by inhibiting mitochondrial fission. The precise mechanism by which mdivi-1 reduces ROS is unclear. Two types of complex I inhibitors have been described: Class A compounds that inhibit complex I and increase ROS levels, and Class B that prevent the formation of ROS even in the presence of Class A inhibitors [49]. It is therefore possible that mdivi-1 behaves like a class B inhibitor, or through a different mechanism altogether.

A key methodological advance of our study was the use of microfluidic chambers to allow independent treatment of axons and cell bodies with vincristine and/or mdivi-1. This method allowed us to demonstrate that mdivi-1 acts locally in the axon to delay vincristine-induced axon degeneration. Furthermore, we observed that axon degeneration was more severe in locally treated axons than in neurons treated globally (Figure 3E vs 3B), suggesting a possible adaptive mechanism by which the cell body signals to the axon to attenuate the injury response.

The rapid and scalable generation of iPSC-derived neurons using transcription factors provides an exciting and highly relevant model in which to test the molecular mechanisms underlying CIPN and chemo brain. The i^3^Neuron model has already been paired with CRISPR-interference (CRISPRi) technology [33, 50], highlighting its utility to identify new druggable pathways and develop novel therapeutic strategies for neurological side effects of chemotherapy. Perhaps most importantly, the human neuron platform helps circumvent potential issues of translatability. Most current knowledge on how chemotherapeutic agents cause axon degeneration derives from research performed in mouse cultured neurons, which may or may not translate well to human.

Using treatment-relevant levels of vincristine in human neurons, we have demonstrated that mitochondrial dysfunction and increase in ROS are early and axon-localized events in vincristine-induced axon degeneration. This degeneration can be pharmacologically mitigated by local treatment of axons with mdivi-1 and other antioxidant agents. Our study reveals that the mitigation of vincristine-induced ROS generation is an exciting novel potential therapeutic target for CIPN and chemo brain which should be further investigated. This work also advances our insight into how clinically relevant levels of vincristine causing mild axon degeneration can lead to the neurological side effects of chemotherapy.

## Methods

### Differentiation of i^3^Neurons

I^3^Neurons were differentiated as previously described. Briefly, i^3^ iPSCs were dissociated using Accutase (Life Technologies, cat. no. A1110501). Cells were plated in Matrigel-coated (1:100 Corning) plates in Neuronal induction media on day 0 (Knockout Dulbecco’s modified Eagle’s medium (DMEM)/F12 medium; Life Technologies Corporation, cat. no. 12660012), 1X N2 supplement (Life Technologies, cat. no. 17502048), 1× GlutaMAX (Thermofisher Scientific, cat. no. 35050061), 1× MEM nonessential amino acids (NEAA) (Thermofisher Scientific, cat. no. 11140050), 10 μM ROCK inhibitor (Y-27632; Selleckchem, cat. no. S1049), and 2 μg/ml doxycycline (Clontech, cat. no. 631311). Neuronal induction media was changed once a day for 2 more days. On day 3 of induction, cells were dissociated using Accutase and plated in dishes coated with poly-L-ornithine (PLO; 0.1 mg/ml; Sigma, cat. no. P3655-10MG). Cells were plated in neuronal maturation media (BrainPhys medium (STEMCELL Technologies, cat. no. 05790), 1× B27 Plus Supplement (ThermoFisher Scientific, cat. no. A3582801), 10 ng/ml BDNF (PeproTech, cat. no. 450-02), 10 ng/ml NT-3 (PeproTech, cat. no. 450-03), 1 mg/ml mouse laminin (Invitrogen, cat. no. 23017015), and 2 μg/ml doxycycline). Half of the neuronal maturation media was removed and replenished with fresh media every 2-3 days.

### Axonal separation and vincristine treatment for mass spectrometry

On day 3 of differentiation, 2 million cells were plated into Falcon^®^ Permeable Support for 6-well plate with 1.0 μm transparent PET membrane (Boyden chambers, cat. No. 353102) in 2 mL of neuronal maturation media. Half media changes were performed every 3-4 days. On day 14 of differentiation, the axonal compartment of the chamber was treated with 5 nM vincristine (Sigma, cat. no. V8879-5MG) or DMSO (Simga, cat. no. 276855) for 4 hours. Axonal and cell body fractions were harvested in ice cold RIPA buffer (Invitrogen, cat. no. 89901) supplemented with protease inhibitors (Sigma, 11836153001). Protein concentrations were calculated using a BCA assay (ThermoFisher, cat. No. 23225). 50 μg of protein was boiled in loading buffer (Invitrogen cat. no. NP0008) and loaded into a 7.5% Mini-PROTEAN^®^ TGX^™^ (Biorad, cat. No. 4561023) for 5 minutes until all protein moved into the gel. Bands were subsequently cut and subjected to mass spectrometry.

### Immunofluorescence, vincristine, and mdivi-1, Glutathione and MitoQ treatments

For immunofluorescent staining of i^3^Neuron axons, cells were plated on a PLO-coated 8-well plate (Ibidi, cat. No. 80827) by adding a 10 μl drop of neuronal maturation media containing ~100,000 cells to the center of each well. Cells were placed in the cell culture incubator for 20 minutes to allow for attachment to the center of the well. 200 μL of neuronal maturation media was then added to the well. For immunofluorescent staining of whole neurons, cells were plated on PLO-coated 96 well plates (PerkinElmer cat. No. 6055300) with 50,000 cells/well. On day 10 or 11 of neuronal differentiation, cells were treated with 5 nM vincristine, 50 μM mdivi-1 (Sigma, M1099), 5mM Glutathione (reduced) (Sigma, G-4251) and 5μM Mitoquinone (MitoQ) mesylate (Selleckchem, S8978). After treatment, cells were fixed in cold 4% PFA for 5-10 minutes. Cells were permeabilized using 0.1% TritonX in PBS for 5 mins and blocked in 5% normal donkey serum (NDS) in PBS at room temperature for 1 hour. Cells were immunostained with anti-beta-3 Tubulin (Invitrogen, cat. no. 2G10-TB3, 1:500) in 2.5% NDS overnight at 4°C. Cells were washed three times with PBS and stained with the appropriate secondary antibody at a concentration of 1:500 in 2.5% NDS for 1 hour at room temperature. Cells were washed three times with PBS and DAPI was used as a nuclear counterstain.

### DHE and ROS quantification

Cells were plated in an 8-well plate and treated with vincristine and mdivi-1 as described above. Cells were treated with DHE (ThermoFisher, cat no. D11347) and MitoSOX^™^ Red Mitochondrial (ThermoFisher, cat. No. M36008). For quantification, 5-8 images of the axons were taken using a Zeiss 800 LSM Scanning Microscope. Fluorescent intensity of DHE and MitoSOX^™^ was quantified using FIJI imaging analysis software and normalized to the intensity area of tubulin (total axonal area) or mitoTracker Green (mitochondrial area) respectively in a blinded manner.

### Mass spectrometry

In-gel samples (~20 μg per sample) were reduced using 10 mM Tris(2-carboxyethyl) phosphine hydrochloride at room temperature for 1 hour and alkylated with 10 mM N-Ethylmaleimide for 10 minutes. Samples were digested with trypsin (Promega) with a trypsin:sample ratio of 1:20 (w/w) at 37°C for 18 hours. Tryptic digests were extracted from the gel and cleaned with an Oasis HLB μElution plate (Waters). Peptides were separated on an ES802A column. Mobile phase B (98% ACN, 1.9% H2O, 0.1% formic acid) amount was increased from 3% to 20% over 83 minutes. Mobile phase A composition is 98% H2O, 1.9% ACN, 0.1% formic acid. LC-MS/MS data were acquired in data-dependent mode. The MS1 scans were performed in orbitrap with a resolution of 120K, a mass range of 400-1500 m/z, and an AGC target of 4 x 10^5^. The quadrupole isolation window is 1.6 m/z. The precursor ion intensity threshold to trigger the MS/MS scan was set at 1 x 10^4^. MS2 scans were conducted in ion trap. Peptides were fragmented with CID method and the collision energy was fixed at 30%. MS1 scan was performed every 3 seconds. As many MS2 scans were acquired within the 3 second cycle.

Proteome Discoverer software version 2.4 was used for protein identification and quantitation. Raw data were searched against Sprot Human database. Up to 2 missed cleavages were allowed for trypsin digestion. NEM on cysteines was set as fixed modification. Mass tolerances for MS1 and MS2 scans were set to 10 ppm and 0.6 Da, respectively. Percolator was used for PSM validation. The search results were filtered by a false discovery rate of 1% at the protein level. Protein abundance values were calculated for all proteins identified by summing the abundance of unique peptides matched to that protein. Protein ratios were calculated by comparing the protein abundances between two conditions with normalization. Adjusted p-values were calculated with the Anova method.

### Seahorse analysis

For Seahorse analysis, 50,000 cells per well were plated after 3 days of induction in a specialized Seahorse analyzer 96-well plate (Agilent, cat. no. 103774-100). 10-12 day old i^3^Neurons were treated with 5 nM vincristine. Oligomycin (2 μM), FCCP (1 μM) and rotenone/antimycin A (500 nM) were added at as shown in figure 2A. Cells were washed in PBS once and incubated in a hypoxic chamber following the manufacturer’s procedure. Seahorse analysis was performed using a Seahorse XFe96 Analyzer (Agilent). The average value for three wells was used for analysis.

### Western blotting

10-14 day old i^3^Neurons were harvested in ice cold RIPA buffer (Invitrogen, cat. no. 89901) supplemented with protease inhibitors (Sigma, 11836153001) and PhosStop (Sigma cat. no. 4906845001). Protein concentrations were calculated using a BCA assay (ThermoFisher, cat. no. 23225). 10-20 μg of total protein lysate was loaded into a 4–20% Mini-PROTEAN^®^ TGX^™^ Precast Protein Gels (Biorad, cat. no. 4561095). Gels were transferred into PVDF membranes and blocked for 1 hour at room temperature in 5% BSA. Membranes were incubated with the appropriate primary antibody in 2.5% BSA solution in a cold room, rocking, overnight. Membranes were washed 3 times with PBS 0.1% Tween solution. Secondary antibody (goat anti-rabbit HRP and IRDye^®^ 680RD Goat anti-Mouse IgG) was incubated for 1 hour at room temperature at a concentration of 1:8000 in 2.5% BSA solution. Membranes were washed 3 times with PBS 0.1% Tween solution prior to developing. Western blots were developed using Clarity Western ECL Substrate (Biorad, cat. no. 1705061) or fluorescence and imaged using a ChemiDoc MP Imaging system (Biorad). Band intensity was quantified using FIJI imaging analysis software. The following antibodies were used at the concentrations stated: anti-DRP1 (Santa Cruz Biotechnology, cat. no. sc-101270, 1:1000), anti-pDRP1 S616 (CST, cat. no. 3455S, 1:1000), anti-β Actin (Sigma, cat. no. A1978, 1:5000), anti-Lamin A (Sigma, cat. no. L1293, 1:1000). Uncropped Western blots are provided in supplementary figure 5.

### Axon degeneration index experiment and quantification

On day 3 of differentiation, i^3^Neurons were dissociated into single cells with Accutase, and plated in Ibidi 8-well μ-Slides (Ibidi, Cat. No. 80826). 50,000 cells were plated in a 5 μL drop of neuronal maturation media in the middle of each well for the axons to grow out from the center. Slides were incubated for 20 minutes at 37°C to allow cells to attach, and 200 μL of neuronal maturation media was then added to each well. Cells were fed with half-media changes every two days until the day of the experiment. Two slides were used for each experiment. On Day 10 or 11 of differentiation, 5 nM vincristine, or a cocktail of 5 nM vincristine and 50 μM mdivi-1, was prepared in neuronal maturation media and added to 6 wells of an 8-well slide. The remaining two wells were fixed at the start of the experiment for the untreated timepoint. Two wells of each slide were fixed at 4 hours, 8 hours, and 24 hours after treatment using 4% cold PFA for 5-10 minutes, then washed three times with PBS. The slides were subsequently blinded for processing and imaging. Cells were immunostained with βIII-Tubulin primary antibody and AlexaFluor 594 secondary antibody and DAPI as described above, and images were acquired on a Zeiss LSM 800 laser-scanning confocal microscope. Five images of axons from two technical replicate wells were taken, for a total of 10 images per timepoint per treatment, for three biological replicates. Images were taken at the distal-most end of the neurons to ensure axons, and not dendrites, were being captured. An axon degeneration index for each image was calculated as follows: 8-bit maximum intensity projections were binarized and thresholder [22] in the FIJI imaging analysis software. The same threshold was used for each image analyzed across biological replicates. The total number of black pixels was measured to be the total axonal area. The degenerated area was measured to be the total area of particles whose pixel circularity was 0.2-1.00. The axon degeneration index (ADI) is calculated as the degenerated area divided by the total axonal area (Sup. Fig. 1).

### Generation and validation of DRP1 knockdown

Control and DRP1 gRNA were cloned into a plasmid containing a U6 promoter using BstXI and BlpI restriction enzymes (NEB) (gifted by Dr. Michael Ward, NINDS). Control gRNA sequence: GGA CTA AGC GCA AGC ACC TA. DNM1L gRNA sequence: GGG AGG AAG GAG GCG AA CTG. Constructs ware verified by Sanger Sequencing and packaged into lentivirus for transduction of iPSCs as follows. In one well of a 6-well plate, 1 million Lenti-X HEK293T cells (Takara Bio, cat. no. 632180) were seeded in 2 mL DMEM (Gibco, cat. no 11995065) supplemented with 10% FBS (Gibco; cat. no. 10437028). The next day, a transfection mix was prepared containing 1 μg DNM1L or Control gRNA-containing plasmid, 3 μg of third generation packaging mix (1:1:1 mix of three plasmids), 12 μL Lipofectamine 3000 Reagent (ThermoFisher, cat. no. L3000008), and 200 μL Opti-MEM I Reduced Serum Medium (GIBCO, cat. no. 31985070). The mix was vortexed, spun down briefly, incubated at room temperature for 20 minutes, then added dropwise to the HEK293T cells and gently swirled to mix. The next day, the media was replaced with 3 ml fresh 10% FBS DMEM supplemented with 1:500 ViralBoost (Alstem, Cat. No. VB100). After two days of incubation, the media was collected into a 15 ml Falcon tube, supplemented with 1 ml Lenti-X Concentrator (Takara Bio; Cat. No. 631231), mixed thoroughly, and stored at 4°C for 48 hours. The supernatant was then spun down at 4°C for 45 minutes at 1,500 x g. The supernatant was aspirated, and the pellet was resuspended in 300 μL of PBS.

CRISPRi-i^3^ iPSCs containing a CAG promoter-driven dCas9-BFP-KRAB cassette inserted into the CLYBL safe harbor locus were the kind gift of Dr. Michael Ward [33]. 500,000 cells were seeded in one well of a Matrigel-coated 6-well plate and transduced with 100 μL of DNM1L, or control sgRNA vector lentivirus in 2 ml mTESR Plus Basal Media with ROCK inhibitor. The next day, fresh media was added containing 1 μg/ml puromycin (Gibco; cat. no. A1113803). Transduced CRISPRi-i^3^N iPSCs were subsequently passaged twice with puromycin until all cells were visibly expressing BFP. One well of a 60-80% confluent 12-well was harvested with 700 μl QIAzxol Lysis Reagent (Qiagen; Cat. No. 79306) for subsequent RNA extraction with RNeasy Mini Kit (Qiagen; Cat. No. 74104) and cDNA synthesis (random hexamers method) with SuperScript III First-Strand Synthesis System (Invitrogen; Cat. No. 18080051) with an input of 1 μg of RNA. Samples were prepared for quantitative real-time PCR in technical duplicates. qPCR was performed with SYBR Green Quantitative RT-qPCR Kit (Sigma-Aldrich; Cat. No. QR0100) using GAPDH as a housekeeping gene, with the CFX Connect Real-Time PCR Detection System (BioRad; Cat. No. 1855201). The following primers were used. GAPDH RNA Fwd: AATGGGCAGCCGTTAGGAAA GAPDH RNA Rev: GCGCCCAATACGACCAAATC DNM1L RNA Fwd: GTTGATCCACTTGGTGGCCT DNM1L RNA Rev: GCCGCTTCACCAGTAACTCA. Expression fold changes were calculated using the ΔΔCt method.

### Annexin V staining

i^3^Neurons were plated in the center of 8-well slides as described above for axons to grow outwards. At 10-14 days old, i^3^Neurons were treated with 5 nM vincristine or 5 nM vincristine with 50 μM mdivi-1. 23 hours after treatment, the cells were incubated with 1:1000 Annexin V, AlexaFluor^TM^ 488 conjugate (Invitrogen; Cat. No. A13201) for 1 hour at 37°C. Images were acquired 24 hours after treatment, in a stage top 5% CO_2_- and 37°C temperature-controlled incubator on a Zeiss LSM 800 laser scanning confocal microscope.

### Generation of SARM1 KO iPSCs

WT iPSCs were transfected with two gRNAs targeting exon 2 of the SARM1 gene gRNA1: AATGCGCGCCACGCGGTCT and gRNA2: CTGTATTGGTGCCGCCGCA. Single cell clones were screened by PCR and sequencing confirmed the deletion of a 161 bp fragment in the SARM1 KO iPSCs.

### Statistical analysis

All quantification of microscopy images was performed blinded. Statistical analyses were performed using GraphPad Prism 9. Gene ontology analysis was performed in R version 4.0.3 (2020-10-10). String analysis was performed with https://string-db.org version 11.5.

## Acknowledgements

We would like to acknowledge the NICHD microscopy facility, with special thanks to Dr. Vincent Schram. We would also like to thank Dr. Yan Li from the NINDS mass spectrometry facility for help generating the proteomics data, and Dr. Alexander Chesler for helpful comments on the manuscript. This work was funded by the Intramural Research Program at the National Institutes of Health (IRP-NIH) through NICHD (ZIA-HD008966 to CELP) and the ALS association (ALSA) Milton Safenowitz Postdoctoral Fellowship (JGD).

## Contributions

Experimental design and conceptualization: JGD, ALS, CELP

Execution and analysis of experiments: JGD, ALS, MN

Writing: JGD, ALS, CELP

## Figure Legends

**Supplementary Figure 1:**
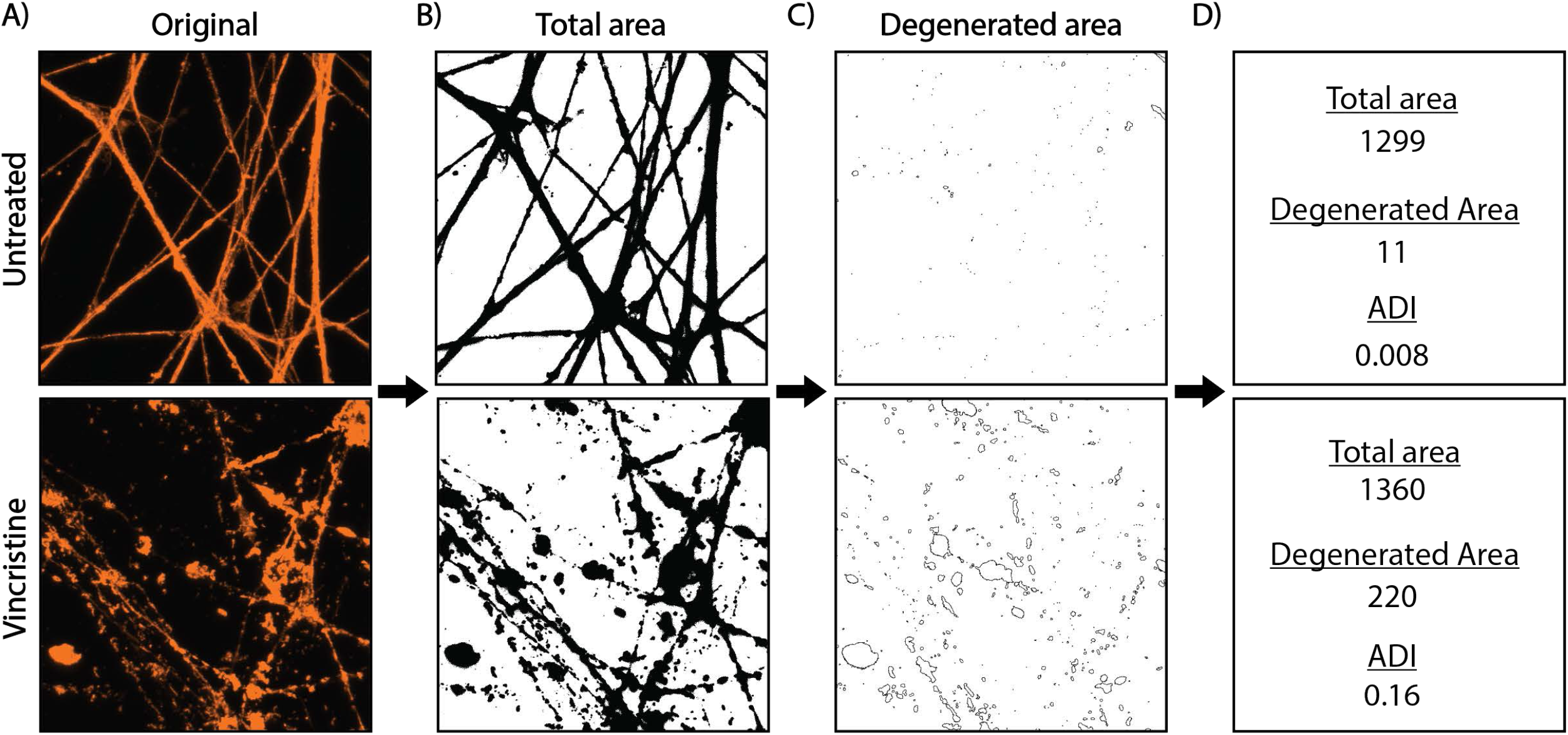
Quantification of axon degeneration index. A) Representative original images of untreated axons (top) and 5 nM vincristine-treated axons (bottom). B) Original images were binarized and adjusted to a determined threshold in ImageJ that was used for all images across biological and technical replicates. Area (px^2^) of the black pixels in the binarized image is considered total axonal area. C) Binarized images are analyzed for particles with pixel units greater than or equal to 2, with a circularity of greater than or equal to 0.2. Total area of these particles is considered the degenerated area (px^2^). D) The axon degeneration index = degenerated area/ total axonal area.

**Supplementary Figure 2:**
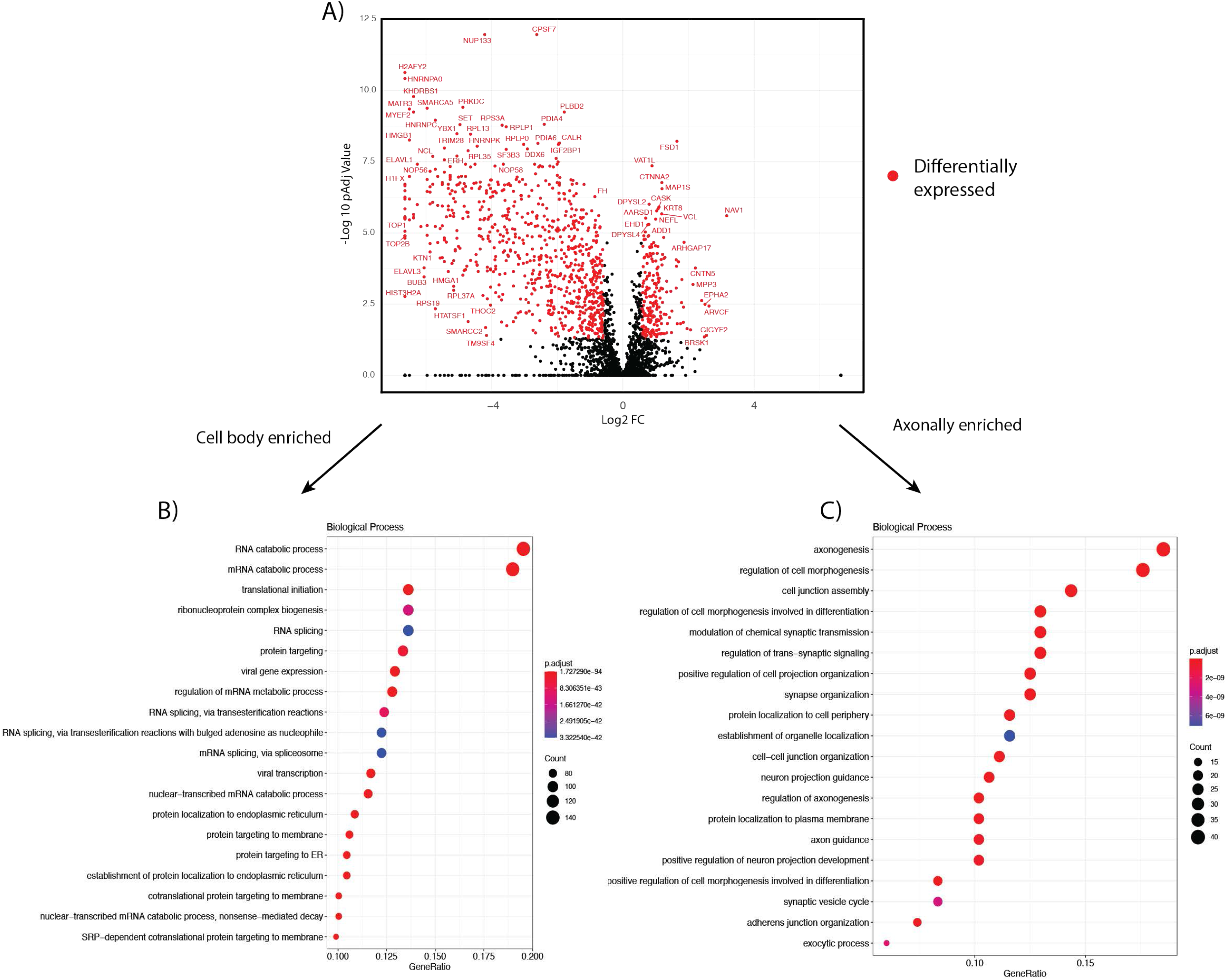
Mass spectrometry of i^3^Neuron cell bodies and axons reveals differences in protein composition. A) Volcano plot showing the significant proteins up- and down-regulated in axons versus cell bodies of i^3^Neurons. (>2 unique peptides detected, Fold Change (FC) >1.5, p. adj. value <0.05.) B) Dot plot showing top 20 enriched biological process gene ontology analysis categories for proteins enriched in the cell body of i^3^Neurons ranked by gene ratio. C) Dot plot showing top 20 enriched biological process gene ontology analysis categories for proteins enriched in the axons of i^3^Neurons ranked by gene ratio.

**Supplementary Figure 3:**
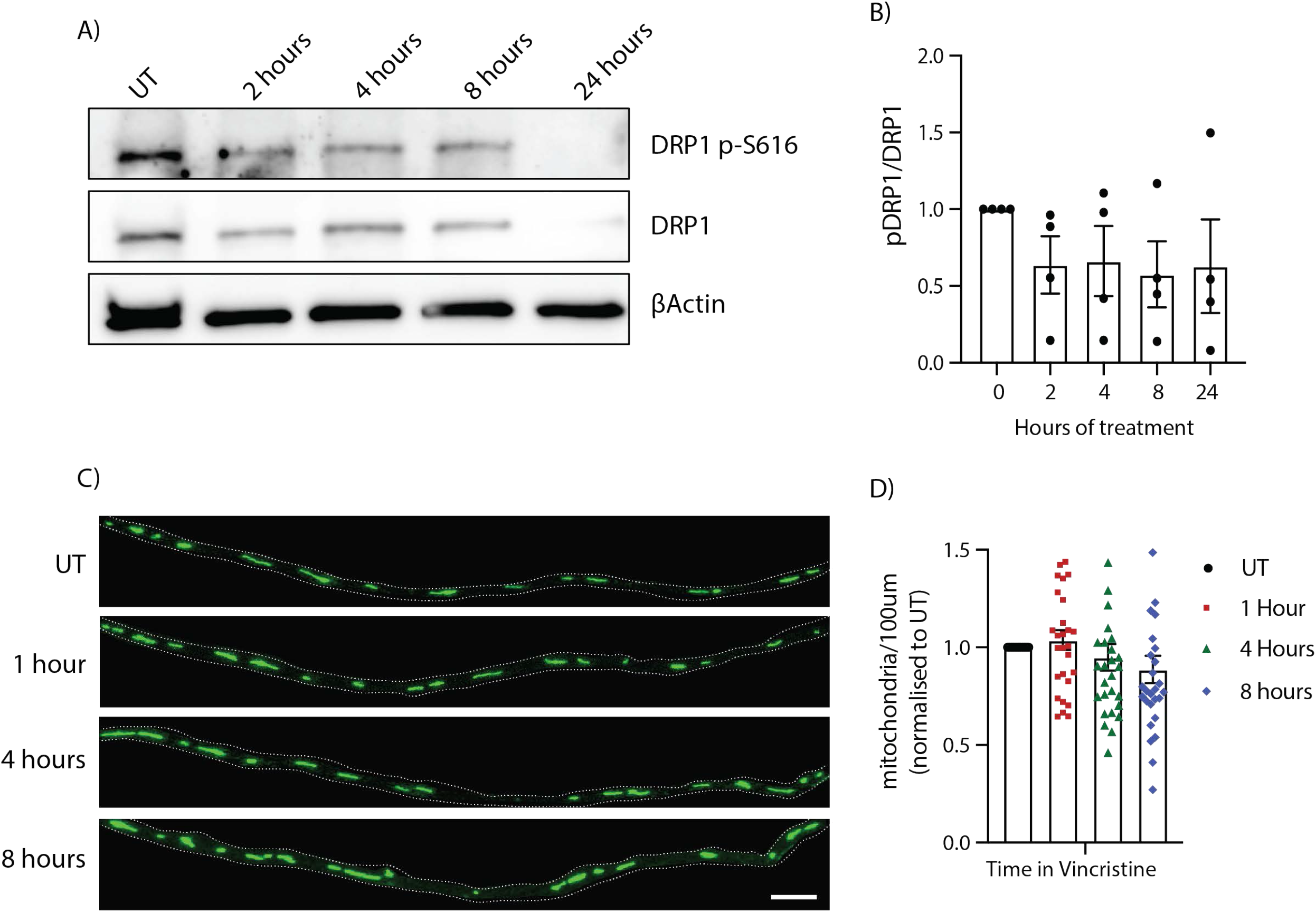
Vincristine does not induce mitochondrial fission in i^3^Neurons. A) Representative western blots of i^3^Neurons treated with 5 nM vincristine for 2, 4, 8, and 24 hours. Immunoblot for p-S616 DRP1, total DRP1 and loading control β actin. Uncropped western blots presented in Sup Fig 5. B) Quantification of pDRP1 (S616)/ total DRP1 levels in 5 nM i^3^Neurons treated with 5 nM vincristine for 2, 4, 8, and 24 hours. Results normalized to UT. Results are represented as mean ± SEM. N=4 independent differentiations. One-way ANOVA. No significant changes observed. C) Representative images of mitoGFP transduced neurons treated with 5 nM vincristine for 0, 2,4 and 8 hours. Dotted line represents the axon. Scale bar = 10 μm. D) Quantification of the number of axonal mitochondrial particles per 100 μm 0, 1, 4 and 8 hours after 5 nM vincristine. N= 28 axons from 3 different differentiations.

**Supplementary Figure 4:**
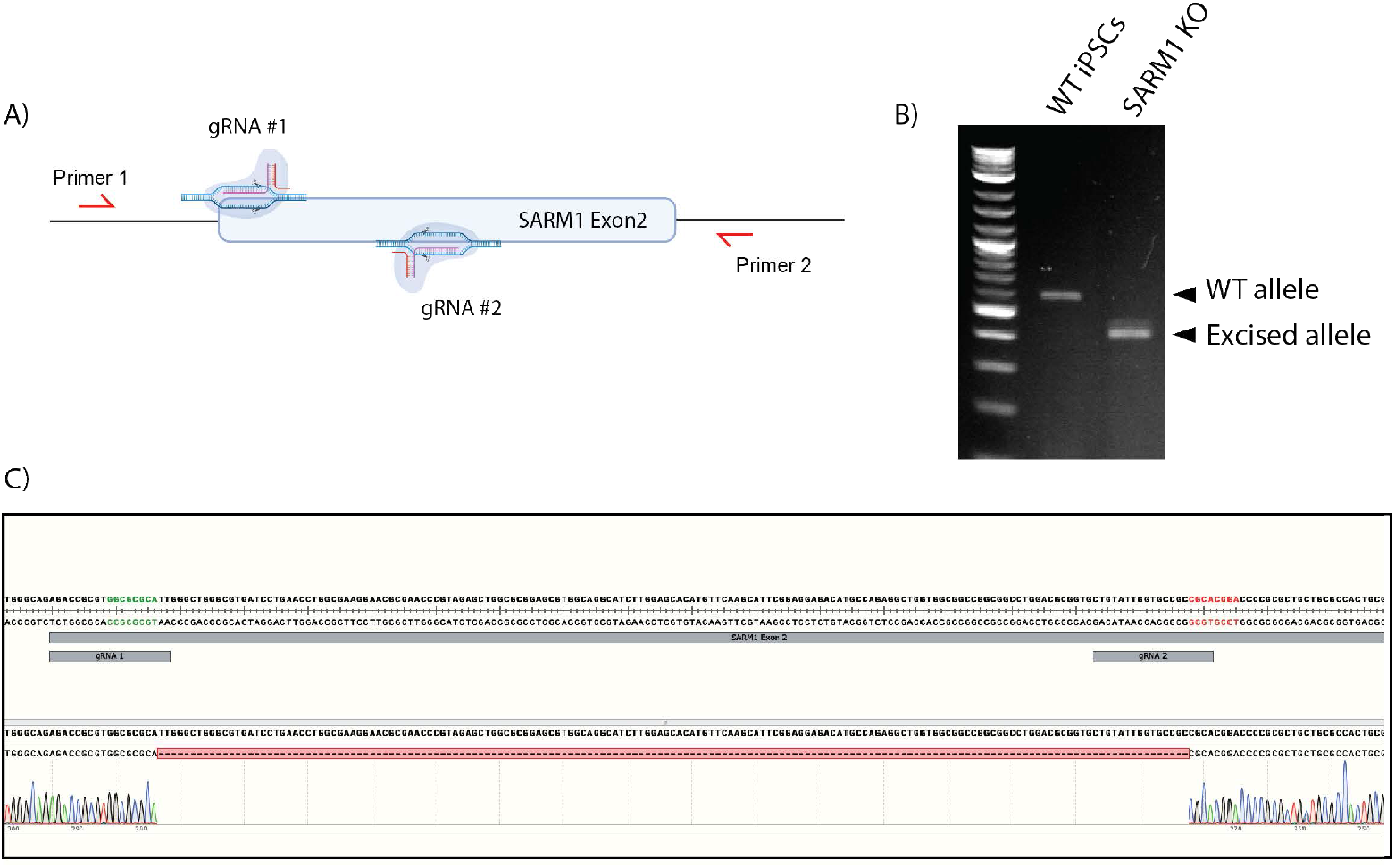
SARM1 knockout generation. A) Schematic representation of SARM1 knockout (KO) generation strategy. WT i^3^ iPSCs were transfected with two gRNAs targeting *SARM1* exon 2. Primers used for knockout validation flanking *SARM1* exon 2 are shown in red. B) PCR of WT and SARM1 KOs using KO primers. SARM1 KOs contain a 161 base pair deletion in exon 2 of the *SARM1* gene, producing a shorter gene product on the agarose gel. C) Sanger sequencing of the *SARM1* deleted allele confirms a 161 base pair deletion resulting in a premature stop codon.

**Supplementary Figure 5:**
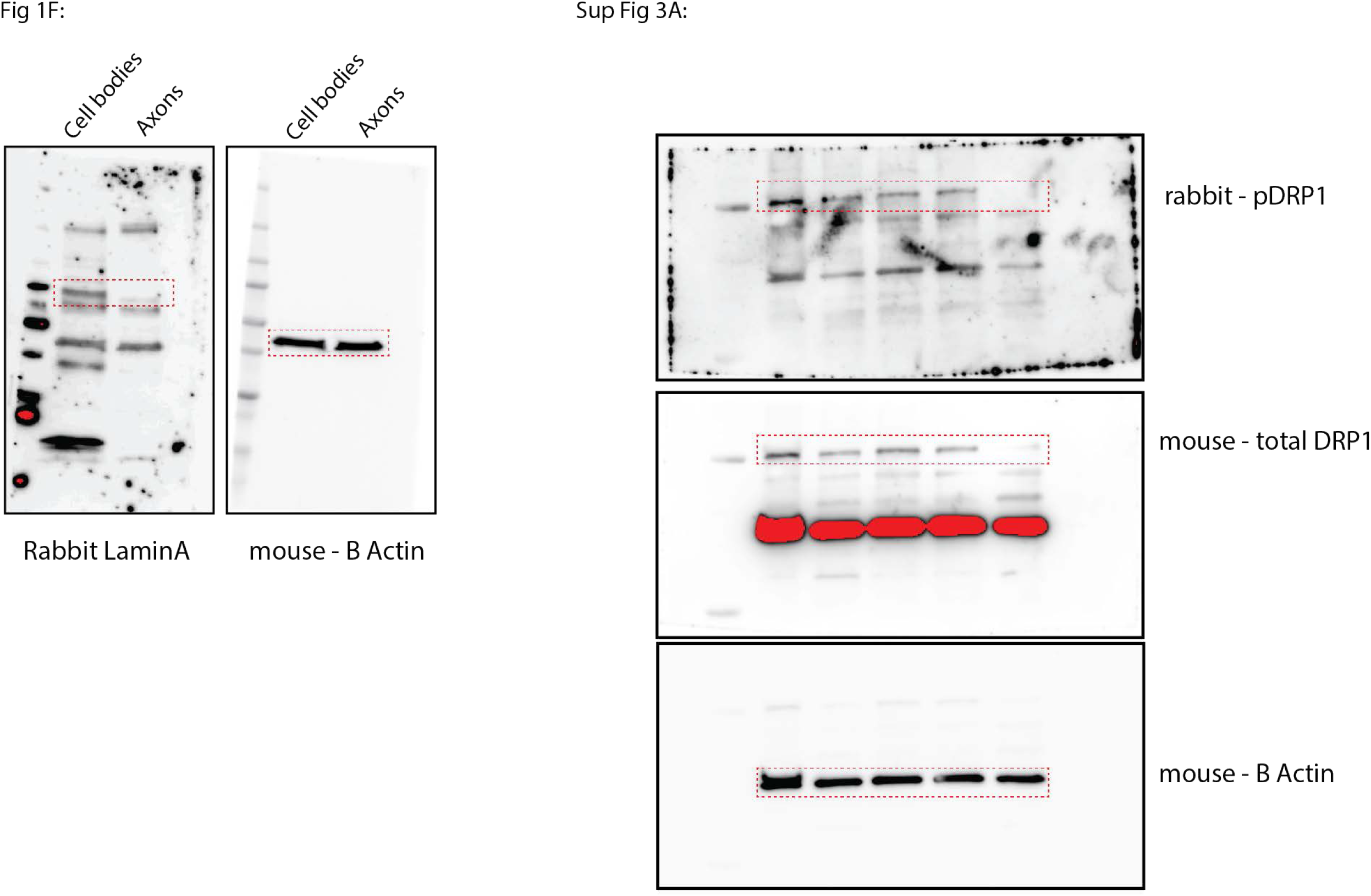
Uncropped Western blots.

## References

1. Geisler, S., Vincristine-and bortezomib-induced neuropathies - from bedside to bench and back. Exp Neurol, 2021. 336: p. 113519.

2. Verstappen, C.C., et al., Dose-related vincristine-induced peripheral neuropathy with unexpected off-therapy worsening. Neurology, 2005. 64(6): p. 1076–7.

3. Diouf, B., et al., Association of an inherited genetic variant with vincristine-related peripheral neuropathy in children with acute lymphoblastic leukemia. JAMA, 2015. 313(8): p. 815–23.

4. Torre, M., et al., Elevated Oxidative Stress and DNA Damage in Cortical Neurons of Chemotherapy Patients. J Neuropathol Exp Neurol, 2021. 80(7): p. 705–712.

5. Melendez, D.M., et al., Spatial memory deficits after vincristine-induced lesions to the dorsal hippocampus. PLoS One, 2020. 15(4): p. e0231941.

6. Zimmer, P., et al., Post-chemotherapy cognitive impairment in patients with B-cell non-Hodgkin lymphoma: a first comprehensive approach to determine cognitive impairments after treatment with rituximab, cyclophosphamide, doxorubicin, vincristine and prednisone or rituximab and bendamustine. Leuk Lymphoma, 2015. 56(2): p. 347–52.

7. Siegel, R.L., K.D. Miller, and A. Jemal, Cancer statistics, 2020. CA Cancer J Clin, 2020. 70(1): p. 7–30.

8. Bates, D. and A. Eastman, Microtubule destabilising agents: far more than just antimitotic anticancer drugs. Br J Clin Pharmacol, 2017. 83(2): p. 255–268.

9. Smith, J.A., et al., Structural Basis for Induction of Peripheral Neuropathy by Microtubule-Targeting Cancer Drugs. Cancer Res, 2016. 76(17): p. 5115–23.

10. Berbusse, G.W., et al., Mitochondrial Dynamics Decrease Prior to Axon Degeneration Induced by Vincristine and are Partially Rescued by Overexpressed cytNmnat1. Front Cell Neurosci, 2016. 10: p. 179.

11. Xu, J., et al., EXPRESS: Methylcobalamin ameliorates neuropathic pain induced by vincristine in rats: Effect on loss of peripheral nerve fibers and imbalance of cytokines in the spinal dorsal horn. Mol Pain, 2016. 12.

12. Chen, X.J., L. Wang, and X.Y. Song, Mitoquinone alleviates vincristine-induced neuropathic pain through inhibiting oxidative stress and apoptosis via the improvement of mitochondrial dysfunction. Biomed Pharmacother, 2020. 125: p. 110003.

13. Geisler, S., et al., Vincristine and bortezomib use distinct upstream mechanisms to activate a common SARM1-dependent axon degeneration program. JCI Insight, 2019. 4(17).

14. Essuman, K., et al., The SARM1 Toll/Interleukin-1 Receptor Domain Possesses Intrinsic NAD(+) Cleavage Activity that Promotes Pathological Axonal Degeneration. Neuron, 2017. 93(6): p. 1334–1343 e5.

15. Diouf, B., et al., Identification of small molecules that mitigate vincristine-induced neurotoxicity while sensitizing leukemia cells to vincristine. Clin Transl Sci, 2021. 14(4): p. 1490–1504.

16. Chen, Y.H., et al., SARM1 is required in human derived sensory neurons for injury-induced and neurotoxic axon degeneration. Exp Neurol, 2021. 339: p. 113636.

17. Fernandopulle, M.S., et al., Transcription Factor-Mediated Differentiation of Human iPSCs into Neurons. Curr Protoc Cell Biol, 2018. 79(1): p. e51.

18. Summers, D.W., A. DiAntonio, and J. Milbrandt, Mitochondrial dysfunction induces Sarm1-dependent cell death in sensory neurons. J Neurosci, 2014. 34(28): p. 9338–50.

19. Sur, M., et al., Sarm1 induction and accompanying inflammatory response mediates age-dependent susceptibility to rotenone-induced neurotoxicity. Cell Death Discov, 2018. 4: p. 114.

20. Miller, B.R., et al., A dual leucine kinase-dependent axon self-destruction program promotes Wallerian degeneration. Nat Neurosci, 2009. 12(4): p. 387–9.

21. Jackson, D.V., Jr., et al., Pharmacokinetics of vincristine in the cerebrospinal fluid of humans. Cancer Res, 1981. 41(4): p. 1466–8.

22. Sasaki, Y., et al., Nicotinamide mononucleotide adenylyl transferase-mediated axonal protection requires enzymatic activity but not increased levels of neuronal nicotinamide adenine dinucleotide. J Neurosci, 2009. 29(17): p. 5525–35.

23. Zhao, R.Z., et al., Mitochondrial electron transport chain, ROS generation and uncoupling (Review). Int J Mol Med, 2019. 44(1): p. 3–15.

24. Dikalov, S.I. and D.G. Harrison, Methods for detection of mitochondrial and cellular reactive oxygen species. Antioxid Redox Signal, 2014. 20(2): p. 372–82.

25. Wu, Q., et al., Mitochondrial division inhibitor 1 (Mdivi-1) offers neuroprotection through diminishing cell death and improving functional outcome in a mouse model of traumatic brain injury. Brain Res, 2016. 1630: p. 134–43.

26. Grohm, J., et al., Inhibition of Drp1 provides neuroprotection in vitro and in vivo. Cell Death Differ, 2012. 19(9): p. 1446–58.

27. Nhu, N.T., et al., Effects of Mdivi-1 on Neural Mitochondrial Dysfunction and Mitochondria-Mediated Apoptosis in Ischemia-Reperfusion Injury After Stroke: A Systematic Review of Preclinical Studies. Front Mol Neurosci, 2021. 14: p. 778569.

28. Bordt, E.A., et al., The Putative Drp1 Inhibitor mdivi-1 Is a Reversible Mitochondrial Complex I Inhibitor that Modulates Reactive Oxygen Species. Dev Cell, 2017. 40(6): p. 583–594 e6.

29. Ruiz, A., E. Alberdi, and C. Matute, Mitochondrial Division Inhibitor 1 (mdivi-1) Protects Neurons against Excitotoxicity through the Modulation of Mitochondrial Function and Intracellular Ca(2+) Signaling. Front Mol Neurosci, 2018. 11: p. 3.

30. van Engeland, M., et al., Annexin V-affinity assay: a review on an apoptosis detection system based on phosphatidylserine exposure. Cytometry, 1998. 31(1): p. 1–9.

31. Bloom, A.J., et al., Constitutively active SARM1 variants that induce neuropathy are enriched in ALS patients. Mol Neurodegener, 2022. 17(1): p. 1.

32. Cassidy-Stone, A., et al., Chemical inhibition of the mitochondrial division dynamin reveals its role in Bax/Bak-dependent mitochondrial outer membrane permeabilization. Dev Cell, 2008. 14(2): p. 193–204.

33. Tian, R., et al., CRISPR Interference-Based Platform for Multimodal Genetic Screens in Human iPSC-Derived Neurons. Neuron, 2019. 104(2): p. 239–255 e12.

34. Knott, A.B., et al., Mitochondrial fragmentation in neurodegeneration. Nat Rev Neurosci, 2008. 9(7): p. 505–18.

35. Sun, E., et al., Docosahexaenoic Acid Alleviates Brain Damage by Promoting Mitophagy in Mice with Ischaemic Stroke. Oxid Med Cell Longev, 2022. 2022: p. 3119649.

36. Bordt, E.A., et al., The Non-Specific Drp1 Inhibitor Mdivi-1 Has Modest Biochemical Antioxidant Activity. Antioxidants (Basel), 2022. 11(3).

37. Meng, T.T., et al., Nicotine Causes Mitochondrial Dynamics Imbalance and Apoptosis Through ROS Mediated Mitophagy Impairment in Cardiomyocytes. Front Physiol, 2021. 12: p. 650055.

38. Lou, M.F., Glutathione and Glutaredoxin in Redox Regulation and Cell Signaling of the Lens. Antioxidants (Basel), 2022. 11(10).

39. Ismail, H., et al., Traumatic Brain Injury: Oxidative Stress and Novel Anti-Oxidants Such as Mitoquinone and Edaravone. Antioxidants (Basel), 2020. 9(10).

40. Schinke, C., et al., Modeling chemotherapy induced neurotoxicity with human induced pluripotent stem cell (iPSC)-derived sensory neurons. Neurobiol Dis, 2021. 155: p. 105391.

41. Spijkers, X.M., et al., A directional 3D neurite outgrowth model for studying motor axon biology and disease. Sci Rep, 2021. 11(1): p. 2080.

42. Ohara, R., et al., Modeling Drug-Induced Neuropathy Using Human iPSCs for Predictive Toxicology. Clin Pharmacol Ther, 2017. 101(6): p. 754–762.

43. Wing, C., et al., Application of stem cell derived neuronal cells to evaluate neurotoxic chemotherapy. Stem Cell Res, 2017. 22: p. 79–88.

44. Wheeler, H.E., et al., Modeling chemotherapeutic neurotoxicity with human induced pluripotent stem cell-derived neuronal cells. PLoS One, 2015. 10(2): p. e0118020.

45. Xiong, C., et al., Human Induced Pluripotent Stem Cell Derived Sensory Neurons are Sensitive to the Neurotoxic Effects of Paclitaxel. Clin Transl Sci, 2021. 14(2): p. 568–581.

46. Kellie, S.J., et al., Cerebrospinal fluid concentrations of vincristine after bolus intravenous dosing: a surrogate marker of brain penetration. Cancer, 2002. 94(6): p. 1815–20.

47. Trecarichi, A. and S.J.L. Flatters, Mitochondrial dysfunction in the pathogenesis of chemotherapy-induced peripheral neuropathy. Int Rev Neurobiol, 2019. 145: p. 83–126.

48. Lopez-Fabuel, I., et al., Complex I assembly into supercomplexes determines differential mitochondrial ROS production in neurons and astrocytes. Proc Natl Acad Sci U S A, 2016. 113(46): p. 13063–13068.

49. Fato, R., et al., Differential effects of mitochondrial Complex I inhibitors on production of reactive oxygen species. Biochim Biophys Acta, 2009. 1787(5): p. 384–92.

50. Tian, R., et al., Genome-wide CRISPRi/a screens in human neurons link lysosomal failure to ferroptosis. Nat Neurosci, 2021. 24(7): p. 1020–1034.

